# Inhibition of p65 NF-κB enhances production of galactose-deficient IgA1 through suppression of *C1GALT1* and SP1 in plasmablast-like cell subpopulations

**DOI:** 10.64898/2026.04.30.721982

**Authors:** Taylor Person, Maggie Phillips, Terri Rice, Stacy Hall, Bruce A. Julian, Dana V. Rizk, Jan Novak, Colin Reily

## Abstract

IgA nephropathy (IgAN) is a common primary glomerulonephritis characterized by glomerular immune-complex deposits with (co)dominant IgA. These deposits are enriched for IgA1 glycoforms with some *O-*glycans deficient in galactose (Gd-IgA1). Circulating Gd-IgA1 is bound by IgG autoantibodies to form immune complexes, some of which deposit in glomeruli. Genomic and immunologic studies indicate involvement of pro-inflammatory signaling pathways in the production of Gd-IgA1 in IgAN. Genomic studies identified multiple genetic loci associated with IgAN and suggested a convergence on the NF-κB pathway, including *RELA*, the gene encoding the NF-κB subunit p65. However, the mechanisms by which NF-κB pathways may affect *O*-glycosylation in IgA1-producing cells are unknown. Using EBV-immortalized B cells derived from peripheral-blood mononuclear cells of IgAN patients and healthy controls that have constitutively activated NF-κB, we report that inhibition of NF-κB/p65 by a selective IKKβ inhibitor TPCA-1 reduced phosphorylation of NF-κB/p65 at S536 and decreased production of IgA1 and, conversely, increased Gd-IgA1 production. This was likely related to reduced expression of *C1GALT1* gene that encodes the enzyme responsible for galactosylation of IgA1 *O*-glycans. Flow-cytometry imaging revealed changes in nuclear translocation and co-localization of the NF-κB/p65 with co-transcriptional factor SP1, a transcriptional activator of *C1GALT1*, suggesting that NF-κB pathway affects IgA1 *O*-glycosylation via SP1 transcriptional control of *C1GALT1* expression. Furthermore, prolonged IKKβ inhibition altered B cell subpopulations, enhancing generation of cells with a plasmablast-like phenotype, characterized by high SSC MFI and CD138 expression. Together, these findings provide functional evidence for involvement of NF-κB/p65 and its transcriptional partners in IgA1 *O*-glycosylation.

**Highlights:**

- IKKβ inhibition reduced *C1GALT1* expression and thereby increased galactose-deficient IgA1 (Gd-IgA1) production in immortalized human B cells.
- SP1^+^ subpopulations, a transcriptional activator of *C1GALT1*, declined after sustained NF-κB inhibition.
- NF-κB inhibition shifted a subpopulation of B cells into a plasmablast-like phenotype.
- This study links NF-κB signaling with the GWAS-identified *RELA* susceptibility locus and IgA1 *O-*glycosylation.

## 1. Introduction

IgA nephropathy (IgAN) is the most common primary glomerulonephritis in many countries [1]. It is diagnosed by kidney biopsy based on detection of IgA as the dominant or co-dominant immunoglobulin in the glomeruli [2, 3]. The IgA is in immune-complex deposits and is of the IgA1 subclass and enriched for glycoforms with some of the hinge-region clustered *O*-glycans lacking galactose (galactose-deficient IgA1; Gd-IgA1) [4, 5]. Complement C3 is usually present. IgAN is associated with significant morbidity and mortality [6, 7] and most IgAN patients progress to kidney failure [8, 9]. Although there are new therapies available and others being tested in clinical trials, there is no cure for this disease [10].

IgA1-containing glomerular immune-complex deposits are thought to originate from the circulation. A multi-hit hypothesis for the pathogenesis of IgAN postulates that in genetically susceptible individuals, Gd-IgA1 is recognized by IgG autoantibodies [11-16]. The resultant immune complexes associate with other proteins, such as complement C3[17, 18]. Some of these circulating complexes enter glomerular mesangium and activate mesangial cells to proliferate and overproduce cytokines, chemokines, and components of extracellular matrix, resulting in kidney injury [19-22]. In this sense, IgAN can be considered an autoimmune disease, with Gd-IgA1 being recognized as an autoantigen [23]. Notably, elevated circulating levels of Gd-IgA1 are associated with disease progression [24, 25]. Therefore, understanding the mechanisms involved in production of Gd-IgA1 is critical for developing new therapies and a cure for this disease.

Core 1 *O*-glycosylation of IgA1 occurs as a step-wise process in the Golgi apparatus of IgA1-producing cells [26]. *N*-acetylgalactosamine (GalNAc) is attached by a GalNAc-transferase to some of the serine or threonine residues in the hinge region, followed by addition of galactose (Gal) by a galactosyltransferase (core 1 synthase, glycoprotein-*N*-acetylgalactosamine 3-β-galactosyltransferase 1; C1GalT1). Production of C1GalT1 protein requires its molecular chaperone C1GalT1C1 (Cosmc) [27, 28]. GalNAc and/or Gal of the core 1 disaccharide can be sialylated. However, premature sialylation of GalNAc by α2,6-sialyltransferase 2 (ST6GalNAc2) prevents addition of Gal. Studies with immortalized IgA1-secreting cell lines derived from peripheral blood of IgAN patients and healthy controls revealed that dysregulated expression and activity of key enzymes are associated with production of Gd-IgA1 [13, 29, 30]. Specifically, reduced expression and activity of C1GalT1 and C1GalT1C1 and elevated expression and activity of ST6GalNAc2 were found in cells derived from IgAN patients compared to the cells derived from healthy controls. Genome-wide association studies (GWAS) found that genetic loci encoding *C1GALT1* and *C1GALT1C1* are associated with serum levels of Gd-IgA1 determined by quantitative lectin ELISA [30]. Gene-specific knock-down of *C1GALT1* and *C1GALT1C1* in IgA1-secreting cell lines confirmed these findings. Furthermore, some cytokines can further dysregulate expression of specific glycosyltransferases and enhance production of Gd-IgA1 by IgA1-secreting cell lines [29, 31, 32]. These cytokines may play a role during synpharyngitic hematuria wherein an upper-respiratory-track infection coincides with disease onset or activity.

GWAS of IgAN patients have identified over 30 genetic risk loci [33-35]. One locus encodes LIF and OSM of the IL-6 family of cytokines [33-35]. LIF can enhance Gd-IgA1 production by IgA1-secreting cell lines derived from IgAN patients due to dysregulated JAK-STAT signaling. Using JAK inhibitors confirmed this pathway [32]. Other genetic loci suggested a convergence on the NF-κB pathway. These GWAS candidates include *REL* and *RELA* genes that encode subunits of the NF-κB transcription factor family, *TNFSF13* gene that encodes A Proliferation-Inducing Ligand (APRIL), and *TNFSF4* and *TNFSF18* genes that encode TNF-related costimulatory ligands [35]. In the canonical NF-κB pathway, TNFα and TLR ligands induce IKKβ phosphorylation of IκBα, releasing p65 (RelA)/p50 dimers to translocate to the nucleus and activate target-gene transcription [36]. The alternative NF-κB pathway utilizes NF-κB inducing kinase (NIK) and is activated by TNF-family cytokines, leading to NIK stabilization, IKKα activation, and p100 processing into p52/RelB heterodimers for gene transcription [37, 38].

As the NF-κB signaling is critical for many B-cell functions, including cellular activation, proliferation, and IgA-class switching, it is difficult to use primary cells to delineate the role of this pathway on a single trait such as *O*-glycosylation of IgA1. Therefore, we used EBV-immortalized B cells that exhibit constitutively activated NF-κB and assessed whether p65 NF-κB inhibition alters IgA1 *O*-glycosylation. Specifically, we used a small-molecule inhibitor of IKKβ, TPCA-1 [39], and tested its effect on production of Gd-IgA1 in immortalized B cells derived from peripheral-blood mononuclear cells (PBMCs) of IgAN patients and healthy controls. We assessed the impact of IKKβ inhibition on IgA1 secretion, degree of IgA1 galactose deficiency, and gene expression of glycosyltransferases involved in IgA1 *O*-glycosylation. Furthermore, we characterized the phenotypic and intracellular signaling consequences of sustained p65 NF-κB suppression across B-cell subpopulations, and connected it to *O*-glycosylation of secreted IgA1.

## 2. Methods

### 2.1 Epstein-Barr Virus-Immortalized B cells

Previously established EBV-immortalized B cells from peripheral-blood mononuclear cells (PBMCs) of healthy controls (12) and IgAN (19) patients were used for these experiments [12, 13, 29]. Laboratory measures at the time of enrollment included serum creatinine (SCr, mg/dL), estimated glomerular filtration rate (eGFR, mL/min/1.73 m^2^), and urine protein-to-creatinine ratio (UPCR, g/g). eGFR was calculated using the 2021 CKD-EPI creatinine equation without a race coefficient (**Supp. Table 1**). Cells were cultured in T75 flasks and maintained with Roswell Park Memorial Institute 1640 media (RPMI1640) supplemented with 10% fetal bovine serum (FBS) and with a cocktail of 1% penicillin and streptomycin. Cells were passaged every two days, and viability and cell number were assessed with Logos Biosystems LUNA-FL™ Dual Fluorescence Cell Counter with AO/PI stain.

### 2.2 Treatment of Cells with IKKβ inhibitor TPCA-1

Cell lines were plated at 1×10^6^ cells/mL in 6-well plates and treated with IKKβ inhibitor TPCA-1 (final concentration of 4 µM in 0.01% DMSO). Vehicle control was 0.01% DMSO. Cells were incubated with TPCA-1 or a vehicle for 1 hr for evaluation of p65 NF-κB inhibition and the effects on glycosyltransferase expression and B-cell signaling. For evaluation of long-term effects of p65 NF-κB inhibition, cells were exposed to TPCA-1 for 48 hr and then assessed by flow cytometry. Cell-culture supernatants were collected after 48 hrs for analysis of IgA1 and Gd-IgA1 concentrations.

### 2.3 IgA1 and Gd-IgA1 Determination

ELISA was used to determine total concentration of secreted IgA1 and Gd-IgA1 from cell lines, as previously described [23, 40]. Briefly, 96-well plates were coated with AffiniPure F(ab’)_2_ fragment of goat IgG specific for human IgA heavy chain (Jackson ImmunoResearch) for IgA capture in both assays. IgA was detected using biotin-labeled F(ab’)_2_ fragment of goat IgG specific for human IgA (GenWay Biotech), followed by HRP-conjugated streptavidin (ExtrAvidin Peroxidase, Sigma). For Gd-IgA1 detection, captured IgA1 was treated with neuraminidase, followed by biotin-conjugated lectin specific for GalNAc from *Helix pomatia* (HPA, Sigma), followed by HRP-conjugated streptavidin (ExtrAvidin Peroxidase, Sigma) and reaction was developed after addition of the substrate OPD and stopped by addition of 5.5% sulfuric acid.

### 2.4 SDS-Polyacrylamide Gel Electrophoresis and Western Blot

Cells were lysed in 50 uL of M-PER buffer (Thermo Fisher Scientific) with 1% 100X halt protease and phosphatase inhibitor cocktail (Thermo Fisher Scientific), and 13 ug protein / lane was loaded into 4-15% PROTEAN TGX stain-free protein gels, 10 well, 30 uL (BioRad). Total protein was separated by SDS-polyacrylamide gel electrophoresis at 95V for 1 hr. Samples were transferred to Immobilon®-FL PVDF membrane (Milllipore-Sigma) for western blot analysis. Antibodies used were anti-phospho-S536-NF-κB (1:1000, Cell Signaling Technology, clone 93H1 Rabbit mAb), NF-κB p65 recombinant rabbit monoclonal antibody (Cell Signaling Technology, clone D14E12, Rabbit mAb). The housekeeping protein used for load controls was glyceraldehyde 3-phosphate dehydrogenase (GAPDH, 1:1000; Li-Cor GAPDH Rabbit Monoclonal, 926-42216). Primary antibodies were detected by secondary antibodies from Li-Cor, IRDye® 680RD Donkey anti-rabbit IgG secondary antibody (1:10,000 for GAPDH, 1:20,000 for NF-κB proteins). PVDF membranes were imaged with the LiCor Odyssey CLx and densitometric evaluation was done with ImageJ software.

### 2.5 Quantitative Real-Time Polymerase Chain Reaction

RNA was isolated from 2×10^6^ cells using RNeasy Plus Mini Kit (Qiagen). Following RNA extraction, complementary DNA (cDNA) synthesis and quantitative PCR amplification were performed in a single reaction using reagents from the Luna Universal One-Step RT-qPCR Kit (New England Biolabs). Reactions were performed in 96-well plates compatible for qPCR. Primers for target genes were generated through Integrated DNA technologies, *C1GALT1* (F-*TCATGCAAGGCATTCAGATG*, R-*ATGGGTTCTTCAGGGTCGTA*), *COSMC* (F-*GCTTTCCTGTCCCCAAGC*, R-*TGCTTTGTCACAGTGTTTGGT*), *ST6GALNAC2* (F-*AAGCTGCTACATCCGGACTTCA*, R-*GGGACAGATCGTGGTTTGCATA*), *IGHA* (F-*CCCCGACCAGCCCCAAGGTCT*, R-*GGCAGGACACTGGAACACGCTGTA*), *JCHAIN* (F-*CCCAGAGCAATATCTGTGATGA*, R-*GGTGGCAGGGAGTTGGTTTAC*). Thermocycling was performed on CFX Opus 96 Real-Time PCR. Results are expressed as fold change vs. the value from the β-actin-encoding housekeeping gene; results were calculated via the 2-ΔΔCt method [41].

### 2.6 Flow Cytometry

Immortalized B cells were fixed and stained for cell-surface targets and then permeabilized (BD Biosciences, BD Cytofix and BD Phosflow Perm Buffer III), followed by staining for intracellular targets **(Supp. Table 2)**. Staining reagents: HPA lectin conjugated with Alexa Fluor 647 (Invitrogen, L32454) (1:100). Phospho-NF-κB p65 (Ser536) monoclonal antibody (T.849.2) (1:1000, Invitrogen, MA5-15160), NF-κB p65 whole protein-recombinant rabbit monoclonal antibody (4-2H22L23, Thermo Fisher Scientific, 701079) (1:1000), goat anti-rabbit IgG(H+L) AF488 (Southern Biotech, 4050-30-8) as secondary antibody for phosphorylated and whole protein p65. SP1-antibody, rabbit monoclonal (Cell Signaling Technology, cat#: 9389S), (1:1000) and AlexaFluor-647 Donkey anti-rabbit IgG (Biolegend, cat #:406414) (1:1000) used for secondary for SP1. DAPI (4’,6-Diamidino-2-Phenylindole, Dilactate) from Thermo Fisher Scientific, (1:1000) was used for nuclear staining. Southern Biotech mouse anti-human IgA PE, OB9130-09 (1:500) was used for detection of cell-surface and total (cell-surface and intracellular) IgA1 in flow cytometry. Side scatter (SSC) was quantified using SSC-A parameters. Imagestream voltage settings were optimized initially using single-color and full-panel-saturation analysis, and then the same settings were applied to all future runs using the same staining workflow. Imagestream files were analyzed using its proprietary Idea’s software. Agilent Opteon was used on the same sample sets analyzed in the Imagestream. Single-color and unstained controls were also used to set up the initial spectral compensation and then used for all sample testing. Gating was performed using FlowJo 10.10. Paired, 2-tailed Student’s t-test was used to compare conditions and subpopulations. Spearman’s rank correlation was used to analyze correlation between flow parameters and Gd-IgA1 production, and p-values were considered significant at < 0.05 using exact permutation distribution.

### 2.7 Clustering, grouping, and module generation

Time-point data, from both 1 hr and 48 hr, were standardized using z-score normalization (52 flow parameters) using the scikit-learner StandardScaler, and hierarchical clustering was performed using Ward linkage, Euclidean distance and the Clustermap package. Grouping was performed using PCA followed by t-SNE (perplexity = 5, random seed = 42, iterations = 2,000). Modules were created using a Spearman correlation matrix between flow parameters with pairwise dissimilarity defined as 1 - |r|, followed by hierarchical clustering using average linkage (cut = 0.1), and applied separately for each timepoint dataset. Every sample and condition had its own module score across the whole set based on the mean z-score of the combined features. Correlation between module and Gd-IgA1 production was assessed using Spearman’s correlation coefficient for each sample module in the set and its corresponding Gd-IgA1 production level.

## 3. Results

### 3.1 TPCA-1 increased Gd-IgA1 production and decreased p65 NF-kB phosphorylation at S536

EBV-immortalized B cells were treated with IKKβ inhibitor TPCA-1. Optimal concentration for NF-κB inhibitor was determined by assessing cell viability and IgA production after 48 hr incubation with different concentrations of the inhibitor **(Supp. Fig. 1)**. Based on these analyses, we used 4 µM TPCA-1, as this concentration did not significantly alter cell growth **(Fig. 1A)** or viability **(Fig. 1B)** compared to control or vehicle-control (equivalent DMSO concentration). B-cell lines treated with TPCA-1 had lower IgA1 production compared to non-treated cells **(Fig. 1C)**. TPCA-1 increased the degree of galactose deficiency in most cell lines tested **(Fig. 1D)**. Furthermore, a relative change in Gd-IgA1 compared to untreated control for each donor showed a significant increase in Gd-IgA1 (p<0.01) **(Fig. 1E)**. We confirmed that TPCA-1 treatment decreased phosphorylation of the canonical p65 NF-κB phosphorylation site at serine 536 (S536) **(Fig. 1F-I)**. After 1 hr of treatment, TPCA-1 significantly reduced the amount of phospho-p65 normalized to GAPDH **(Fig. 1G**, p<0.01). Total p65 NF-κB protein remained unchanged **(Fig. 1H)**, confirming that TPCA-1 did not alter total amount of p65 protein. When the amount of phosho-p65 (S536) was normalized to total p65 NF-κB protein it confirmed a significant reduction of phosphorylation **(Fig. 1I)**.

**Figure 1:**
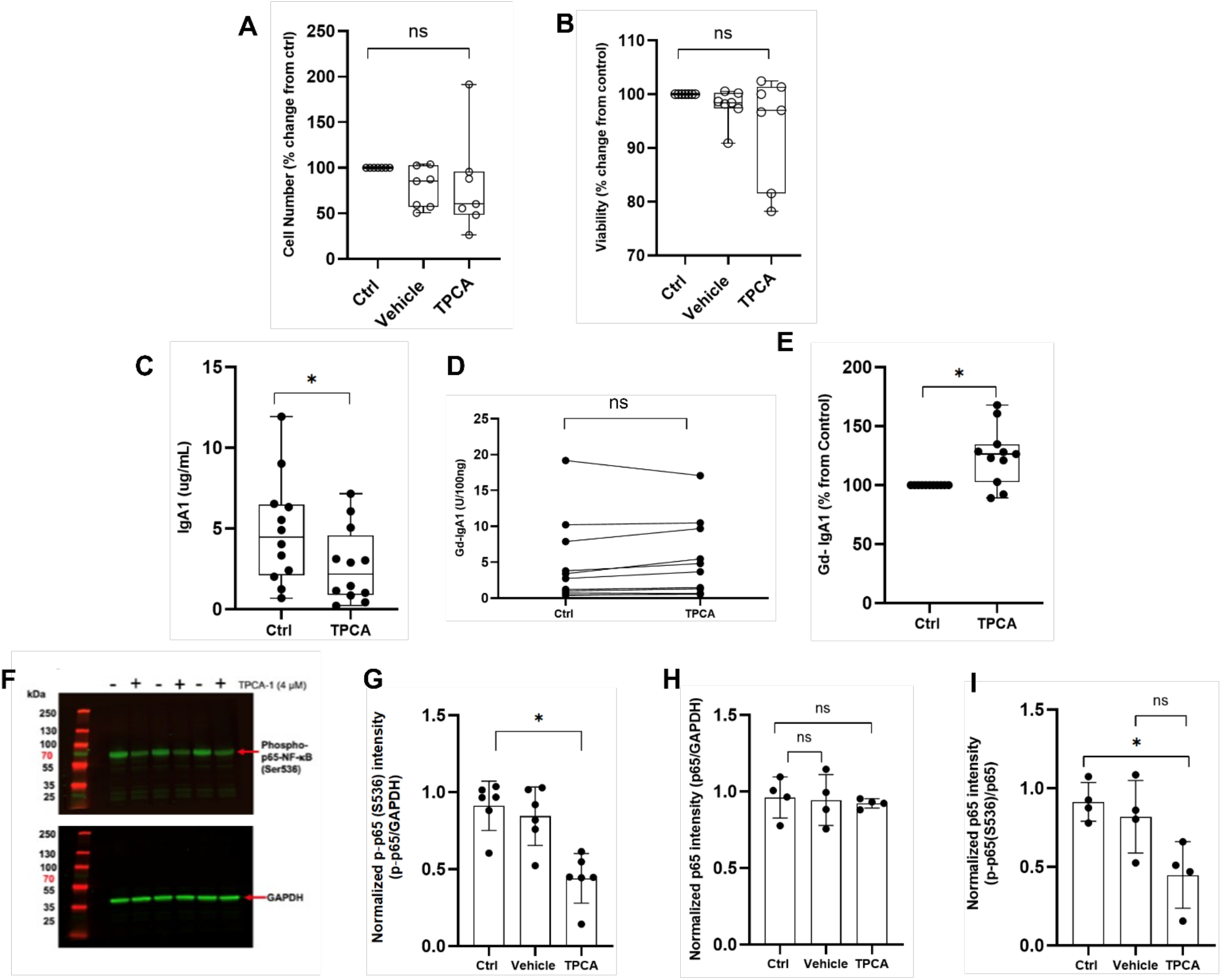
Small-molecule inhibitor of IKKβ TPCA-1 reduced galactosylation of IgA1 and decreased IgA1 secretion without impacting cell proliferation and viability. **A)** Cell number as a % change vs control after TPCA or vehicle treatment for 48 hr, no significant difference. **(B)** Viability of cells as a % change vs control after 48 hr treatment with TPCA or vehicle, no significant difference. **C)** Amount of IgA1 secreted in the cell-culture medium compared to control (Ctrl). Samples were analyzed after 48 hr. **D)** TPCA increased the degree of IgA1 galactose deficiency in most cell lines after 48 hr. **E)** TCPA increased Gd-IgA1 when expressed as a relative change compared to untreated control for each donor. **F)** Representative western blot of phospho-S536 p65 NF-κB (top) and GAPDH (bottom) after 1 hr of treatment. **G)** Densitometric analysis of phospho-S536 p65 NF-κB normalized to GAPDH after 1 hr of treatment. **H)** Densitometric analysis of total p65 NF-κB protein to GAPDH after 1 hr of treatment. **I)** Densitometry analysis of phospho-S536 p65 NF-κB normalized to total p65 NF-κB protein after 1 hr of treatment. *p<0.01. ns = no significance. P values were calculated using Students’ t-test, paired, 2-tailed. n = 6 (A,B,G-I), n = 11 (C-E).

### 3.2 TPCA-1 decreased expression of β1,3-galactosyltransferase-encoding gene C1GALT1

qRT-PCR was used to determine the effects of p65 NF-κB inhibition on expression of genes encoding critical *O-*glycosyltransferases and IgA1 component chains (n=11). These genes included those encoding the key galactosyltransferase, *C1GALT1*, its chaperone *COSMC*, the sialyltransferase *ST6GALNAC2*, and the constant segments of IgA1 heavy chain and J chain, *IGHA1* and *JCHAIN*. J chain is required for production of polymeric IgA. After 1 hr of TPCA-1 treatment, expression of *C1GALT1* decreased in all cell lines **(Fig. 2A**, p=0.01). Expression of *COSMC* and other tested genes did not significantly change **(Fig. 2A,B)**.

**Figure 2:**
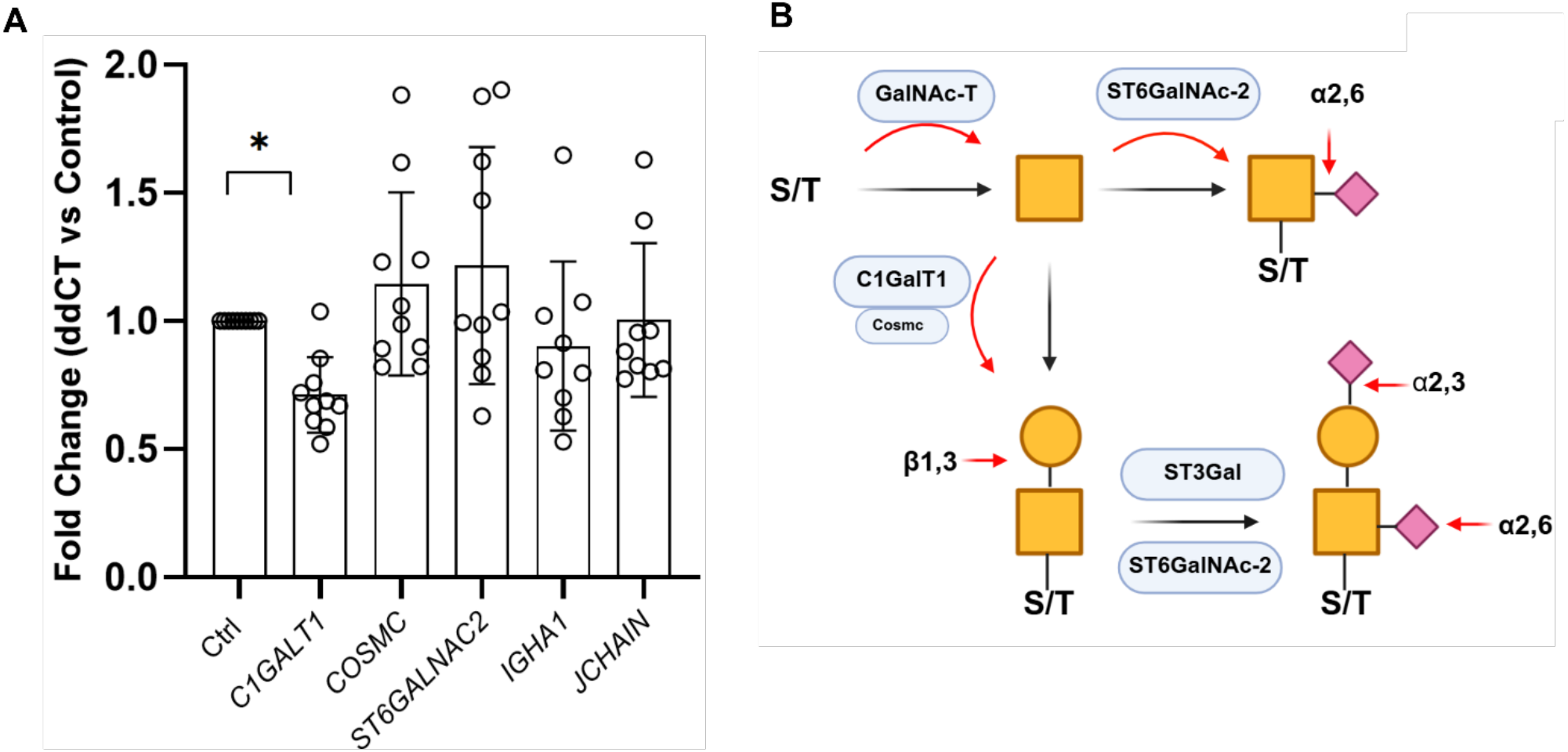
p65 NF-*k*B inhibition at S536 decreased expression of *C1GALT1* but not of other glycosyltransferases or IgA heavy chain or J chain. **A)** B cells treated with TPCA-1 for 1 hr, expression compared to control (normalized to *β-actin*, fold change vs. Ctrl). **B)** Scheme showing core-1 *O*-glycosylation of the IgA1 hinge-region. GalNAc is attached to a serine or threonine residue(s) by a GalNAc transferase enzyme. Usually, GalNAc is next modified by C1GalT1 that adds β1,3-linked Gal; production of the active C1GalT1 requires its chaperone protein Core 1 β1-3-galactosyltransferase-specific chaperone 1 (Cosmc). This structure can then be sialylated, either via α2,6 linkage by ST6GalNAc-2 and/or via α2,3 linkage by ST3Gal. GalNAc can also be prematurely sialylated via ST6GalNAc-2, preventing any other modification, n=9. *p= 0.01. Student’s t-Test, paired, 2-tailed.

### 3.3 TPCA-1 increased the proportion of SSC-high subpopulations which contain lower amounts of p65 NF-kB and SP1

We assessed the phenotypic effects of p65 NF-κB inhibition on B-cell populations by flow cytometry following 1 hr and 48 hr of TPCA-1 treatment. TPCA-1-treated cells demonstrated a rightward shift in SSC-A distribution relative to vehicle-treated and untreated controls **(Fig. 3A)**, corresponding to a significant increase in the frequency of SSC-high cells within the parent population only at 48 hr **(Fig 3B)**. Cells with higher HPA lectin binding, that corresponds to a higher amount of accessible terminal GalNAc on cell-surface glycoconjugates, had a higher SSC-A average value **(Fig. 3C**, p<0.01) than the cells with lower HPA lectin binding. TPCA-1 treatment decreased the proportion of cells with high levels of intracellular p65 NF-κB at the 48-hr timepoint and, conversely, increased the frequency of the population with low levels of intracellular p65 NF-κB at the same timepoint (**Fig. 3D**, p = 0.01). We observed a similar pattern for SP1^+/-^ cells, with no differences in expression detected at 1 hr. After 48 hr, TPCA-1 treatment significantly decreased the frequency of SP1^+^ cells and increased the frequency of SP1^+^ cells (**Fig. 3E**, p = 0.01).

**Figure 3:**
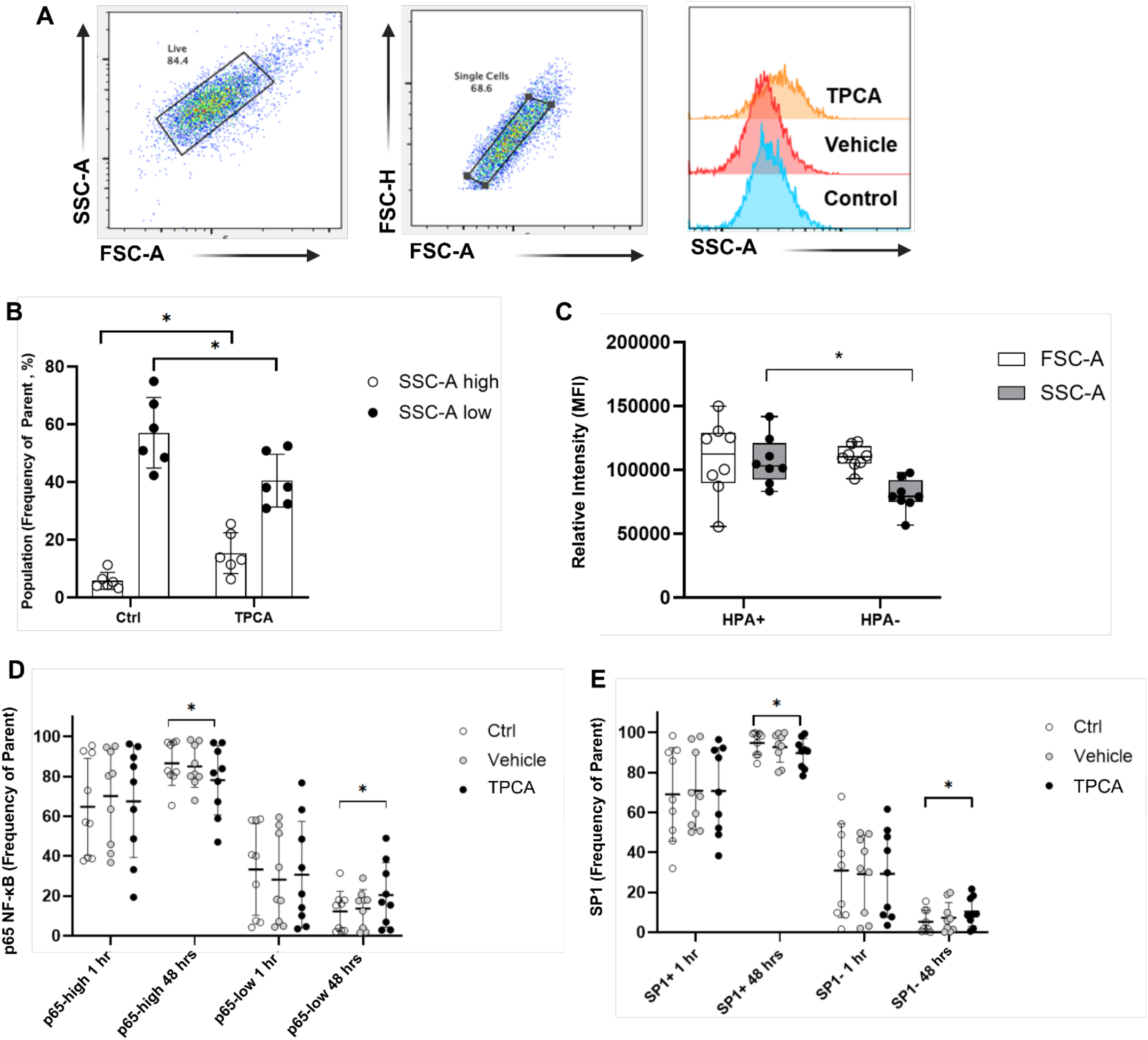
TPCA-1 shifts B cell population to higher SSC and HPA+ phenotype. **A)** B cells exposed to TPCA for 48 hr had increased frequency of populations with higher SSC-A as shown by the right shift. **B)** B cells exposed to TPCA-1 for 48 hr, gated for top 20% and bottom 20% SSC-A populations based on control. **C)** MFI of SSC-A and FSC-A in HPA^+^ and HPA^-^ subpopulations. **D)** Subpopulations of p65 NF-κB gated for top 25% and bottom 25% based on control, grouped by treatment after 48 hr. **E)** B cells exposed to TPCA or vehicle for 48 hr and gated into SP1^+^ and SP1^-^ subpopulations. bar = mean and error bars are +/-SEM; brackets represent paired t-test vs controls. p<0.05 only. n=6 for Figs. 3A,B,C, n=9 for Figs. 3D,E. *p = 0.01. Student’s t-test, paired, 2-tailed.

### 3.4 SSC separates cells into distinct subpopulations with distinct transcriptional states

We determined whether SSC-A low and high subpopulations had any correlation with intracellular signaling components across treatment conditions at 1 hr and 48 hr. We compared average levels of p65 NF-κB, IgA1, and SP1 between SSC-high and SSC-low cells. SSC-low cells trended toward a higher mean fluorescence intensity (MFI) for p65 NF-κB than SSC-high cells at 1 hr, although this difference did not reach statistical significance in any treatment group. At 48 hr, this trend persisted and reached significance in the TPCA-1-treated condition between subpopulations. Total IgA, which reflects cell-surface and intracellular IgA, was significantly higher in SSC-low cells compared to SSC-high cells across all conditions at both timepoints **(Fig. 4B)**. SP1 protein levels at 1 hr in SSC-low cells were significantly higher than in SSC-high cells (**Fig. 4C**, p < 0.01). After 48 hr with TPCA-1 treatment, SP1 expression differences were seen in only TPCA-1-treated cells (p < 0.001); results for the control cells and vehicle-treated cells did not significantly differ from those for the SSC populations, although directional trends toward higher SP1 expression in SSC-low cells remained.

**Figure 4:**
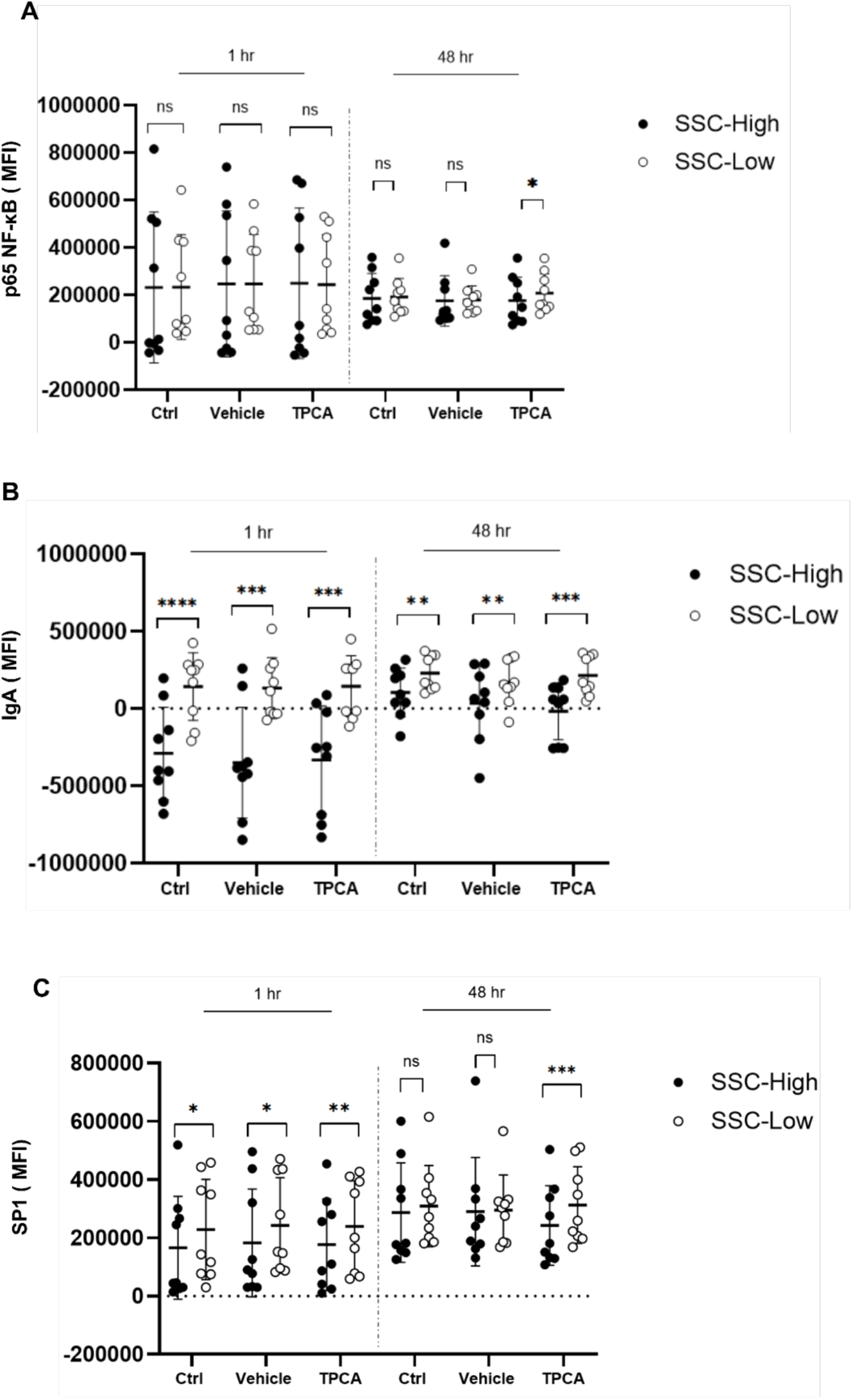
SSC-A separates cells into distinct subpopulations with differential transcriptional controls. B cells treated with vehicle or TPCA-1 for 1 hr or 48 hr and gated into SSC-high and SSC-low populations. **A)** B cells treated for 1 hr or 48 hrs with TPCA-1 or vehicle, gated for SSC-low or high subpopulations and p65 NF-κB total protein MFI was assessed. **B)** B cells treated for 1 hr or 48 hr with TPCA-1 or vehicle, gated for SSC-A low or high subpopulations and IgA (cell surface + intracellular) MFI was assessed. **C)** B cells treated for 1 hr or 48 hr with TPCA-1 or vehicle, gated for SSC-A low or high subpopulations and SP1 total protein MFI was assessed. n=9. *p=<0.05; **p=<0.01; ***p=<0.001; ****p=<0.0001. Student’s t-test paired, 2-tailed.

### 3.5 SSC and IgA^+^ subpopulations exhibit differential nuclear translocation activities of p65 NF-kB and SP1

We determined nuclear localization of the two transcription factors in SSC populations using the ImageStream X. The ImageStream allows visualization of intracellular targets; we used nuclear staining to assess translocation and colocalization movement of our targets **(Fig 5A)**. The similarity score is a quantitative metric within the ImageStream IDEAS software that measures spatial pixel-intensity correlation between two fluorescent targets within a single cell. A higher positive score indicates stronger colocalization. Progressively negative scores indicate weaker colocalization. SP1 nuclear translocation was significantly higher in SSC-low compared to SSC-high cells at 1 hr and at 48 hr **(Fig 5B)**. There was no significant difference in p65 NF-*k*B nuclear translocation between SSC-high and SSC-low populations across any condition or timepoint (data not shown). Because p65 NF-*k*B and SP1 can form a co-transcription complex, we examined the spatial colocalization of p65 NF-*k*B and SP1 protein (CH02 and CH11 in **Fig. 5A**). SSC-low cells had significantly higher p65 NF-*k*B-SP1 colocalization than did SSC-high cells across treatment conditions **(Fig. 5C)**. At 48 hr, the differences between SSC-high and - low populations were no longer significant. P65 NF-κB nuclear translocation was significantly higher in IgA^+^ cells compared to IgA^+^ cells at 1 hr in all groups **(Fig. 5D)**. There was no significance between populations after 48 hr. IgA^+^ cells exhibited significantly higher SP1 nuclear translocation across all conditions at 1 hr and 48 hr across all treatment conditions compared to IgA^+^ cells **(Fig. 5E)**.

**Figure 5:**
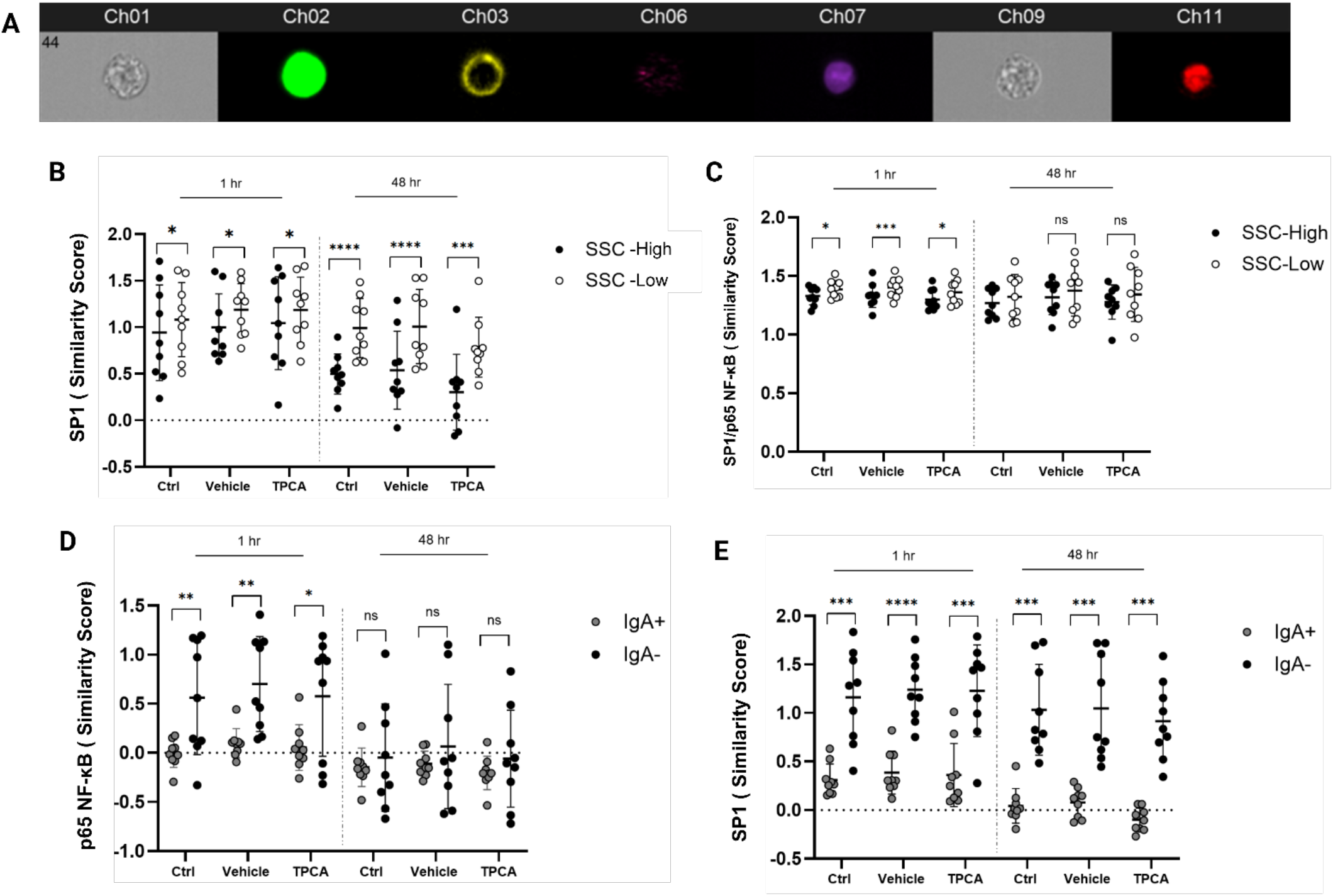
SSC-A and IgA subpopulations exhibit differential nuclear translocation activities of p65 NF-*k*B and SP1. B cells treated with vehicle or TPCA-1 for 1 hr or 48 hr, and split into high and low SSC-A populations or IgA^+^ and IgA^-^ populations. **A)** Sample image of a stained B cell from the ImageStream Ideas software. Ch01/Ch09 brightfield, Ch02 p65 NF-*k*B total protein, Ch06 SSC, Ch07 DAPI, Ch11 SP1 total protein. **B)** SP1 nuclear translocation in high and low SSC-A populations after 1hr and 48 hr of TPCA-1 or vehicle treatment. **C)** Overlapping colocalization of p65 NF-*k*B and SP1 protein at 1hr and 48 hr after TPCA-1 or vehicle treatment. **D)**p65 NF-*k*B nuclear translocation in IgA^+/-^ subpopulations at 1 hr and 48 hr of TPCA-1 or vehicle treatment. **E)** SP1 nuclear translocation in IgA^+/-^ subpopulations at 1 hr and 48 hr of TPCA-1 or vehicle treatment. *p=<0.05;**p=<0.01;***p=<0.001;****p=<0.0001.Student’s t-test paired, 2 -tailed. n=9.

### 3.6 Unsupervised grouping reveals correlation between Gd-IgA1 production and SP1 subpopulations at 48 hr

We performed a Spearman’s correlation analysis of 52 flow parameters against Gd-IgA1 production. Each module was scored using the mean z-score of its member features, and 1 hr and 48 hr timepoints were analyzed independently. At 1 hr, module scores showed non-significant correlations with Gd-IgA1 across all treatment conditions **(Fig. 6A)**, and only 1 of 52 individual flow parameters reached significance **(Fig. 6D)**. At 48 hr, greater differentiation among modules emerged **(Fig. 6B)**; 3 of 52 individual parameters reached significance, all with positive Spearman’s correlation coefficients **(Fig. 6E)**. When module-level correlations with Gd-IgA1 were compared across both timepoints, only one module, M7, reached statistical significance, and only at 48 hr (**Fig. 6C)**. The individual features comprising M7 revealed that its significant members were SP1 MFI measurements within IgA^+^ and SP1^+^ gated subsets **(Fig. 6F)**. The M7 module score at 48 hr positively correlated with Gd-IgA1 production across all the conditions (ρ = 0.48, p = 0.043, **Fig. 6G**). However, M7 module scores did not differ appreciably between control, vehicle-treated, and TPCA-1-treated conditions **(Fig. 6H)**, indicating that M7 module reflects a cell-line-intrinsic property rather than a treatment-responsive change.

**Figure 6:**
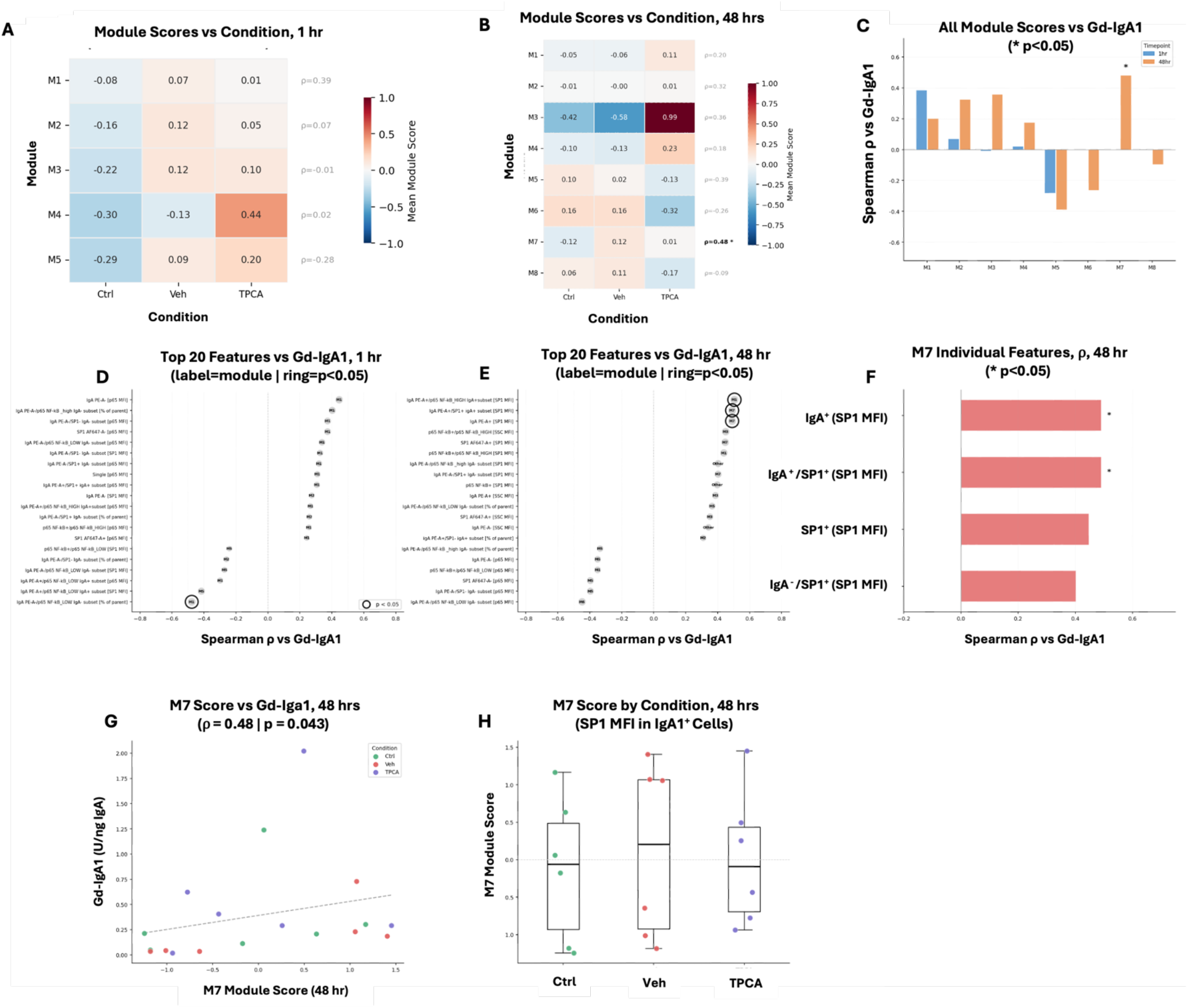
Unsupervised grouping reveals correlation between Gd-IgA1 production and SP1 MFI at 48 hr. Spearman’s correlation analysis of flow parameters to Gd-IgA1 production levels to create a dissimilarity matrix, followed by hierarchical clustering and scored using the mean z-score of its member features. **A)** Heatmap of module scores vs treatment at 1 hr. **B)** Heatmap of module scores vs treatment at 48 hr. **C)** Spearmen score for each module at both timepoints for Gd-IgA1 correlation. **D)** Top 20 flow parameters correlated to Gd-IgA1 for each module at 1 hr. **E)** Top 20 flow parameters correlated to Gd-IgA1 for each module at 48 hr. **F)** Individual Spearmen coefficient flow parameters in the significant M7 module at 48 hr. **G)** Spearmen correlation between individual M7 module scores for each sample condition and Gd-IgA1 at 48 hr. **H)** Chart of M7 module scores for each sample broken down by treatment at 48 hr. Spearman’s correlation analysis, 2-sided, n=6.

## 4 Discussion

The association between an upper-respiratory-tract infection with its inflammation and production of Gd-IgA1 and IgA1 in IgAN is well known, but the mechanisms have not been fully defined. The association of synpharyngitic hematuria with disease onset or clinical flare implicates mucosal inflammation in the pathogenesis of IgAN [42, 43]. Additionally, GWAS studies have identified multiple susceptibility loci involving inflammatory signaling pathways, namely those commonly active during infection, such as IL-6 and the JAK/STAT pathways [31, 32]. Corticosteroid treatments have been used for IgAN patients who are at risk of disease progression. Such therapy often reduces circulatory levels of Gd-IgA1 [44-46]. In our model of EBV-immortalized B cells from patients with IgAN and healthy controls, TPCA-1 increased Gd-IgA1 production in almost all cells (Fig. 1). Notably, we previously found that IL-6-family cytokines increased Gd-IgA1 production, but only in cell lines derived from PBMCs from IgAN patients [29, 31, 32]. The present study used EBV-immortalized cells that were not subcloned, thus retaining the full phenotypic heterogeneity of the donor B-cell populations that survived immortalization. The observation that TPCA-1 consistently increased Gd-IgA1 production irrespective of the donor group suggests that the effect of NF-κB inhibition plays a broad role in autoantigen production. This finding additionally suggests that the relationship between NF-κB signaling and IgA1 glycosylation may not be linear and may involve other signaling components.

Multiple connections between NF-κB signaling pathways and the pathogenesis of IgAN have been published. A 2023 GWAS study identified multiple associated genes converging on this signaling pathway, including *REL* and *RELA* genes that encode subunits of the NF-κB transcription factor pathway, *TNFSF13* gene that encodes A Proliferation Inducing Ligand (APRIL), and *TNFSF4* and *TNFSF18* genes that encode TNF-related costimulatory ligands [35].

p65 NF-κB is not a transcription factor that can bind within the promoter regions of *C1GALT1* or *COSMC*; therefore, transcriptional co-factors may be involved in facilitating p65 NF-κB regulation of these genes. SP1 is a known transcription factor that positively regulates *C1GALT1*, and also forms complexes with p65 NF-κB, making it a prime candidate for investigation in the context of Gd-IgA1 production [47]. Specifically, ChIP analysis and dual-luciferase reporter assays have demonstrated that SP1 directly binds two sites within the *C1GALT1* promoter, and is required for transcriptional activation; mutation of both SP1 binding sites abolishes promoter activity [48]. Our results are consistent with this potential regulatory relationship; TPCA-1 decreased *C1GALT1* expression within 1 hr of treatment (**Fig. 2A**), consistent with potential disruption of transcriptional support at *C1GALT1* promoter. We did not find changes in SP1 or p65 NF-κB after 1 hr of TPCA-1 treatment, but rather after 48 hr, SP1^+^ population was reduced. A significant decrease in the p65-NF-κB high population paralleled the decrease in SP1^+^ population (**Fig. 3D,E**). There is evidence that suggests that SP1 is recruited to its natural promoter sites in a p65 NF-κB-dependent manner [49]. p65 NF-κB/SP1 interactions have been shown to be either activating or repressive, depending on promoter context and presence of other co-regulatory factors [50, 51]. Direct confirmation through ChIP Seq analysis of SP1 and p65 presence at the *C1GALT1* promoter in IgA1-producing B cells will be an important future direction to confirm their respective roles.

The mechanisms regulating Gd-IgA1 production appear to be independent of those controlling total IgA1 output. Baseline IgA1 production levels strongly correlated across treatment conditions, indicating that the determinants of total IgA1 secretion are linearly affected by IKKβ inhibition (**Supp. Fig. 4**). However, there was no correlation between baseline and TPCA-1 treated Gd-IgA1 levels, which suggests p65 NF-κB inhibition has different magnitudes of effect on *O-*glycosylation between cell lines (**Supp. Fig. 4**, p = 0.03, r = 0.957). Together, these findings demonstrate that TPCA-1 may specifically disrupt the glycosylation of IgA1 in a manner independent of IgA1 production. Prior observations support the dissociation between production of IgA1 and its *O*-glycosylation [29, 31]. Some cytokines dysregulated glycosyltransferase expression and increased Gd-IgA1 production (in cells derived from IgAN patients) irrespective of the changes in production of total IgA1 (in cells derived from IgAN patients and healthy controls), showing that these processes are governed by separable regulatory mechanisms [29, 32]. In our work, we observed that *IGHA1* expression was not altered by TPCA-1 treatment at 1 hr **(Fig. 2a)** and total IgA1 MFI (extracellular + intracellular staining) was not significantly altered at 48 hr **(Fig. 4b)**, a surprising finding given the decrease in IgA1 secretion. This result indicates that perhaps transcriptional changes are occurring after TPCA-1 exposure, and that secretion of IgA1 may be independent of glycosylation of IgA1 and cell-surface glycoconjugates. IgA1 *O-*glycosylation occurs post-translationally in the Golgi apparatus; thus, changes in specific glycosyltransferase activity can alter glycosylation without changing production of the immunoglobulin.

TPCA-1 shifted cellular distribution towards cells with higher granularity (more intracellular organelles), indicated by an increase in SSC-A subpopulations; these cells were also enriched for HPA-binding cell-surface glycoconjugates and, thus, elevated amounts of terminal GalNAc **(Fig. 3B,C)**. This finding suggests an increase in a subpopulation of cells associated with Gd-IgA1 production and elevated content of glycoconjugates with terminal GalNAc. Reduction of total IgA1 may be attributed to inhibition of cell proliferation, as there was an insignificant trend toward fewer cells at 48 hr (**Fig. 1A**), rather than suppression of transcriptional production of IgA1, as *IGHA1* expression was unchanged at 1 hr. The increase in Gd-IgA1 production could reflect a mechanistic disruption of the C1GalT1/SP1 axis, which acts independently of total IgA1 production. Thus, TPCA-1 reduced IgA1 production through its effects on cell proliferation while also increasing aberrant *O-*glycosylation of IgA1 through decreased *C1GALT1* expression. Notably, the cells that have increased SSC-A are only a small population of cells. This finding, combined with the increased Gd-IgA1 production, is consistent with evidence that aberrant IgA1 glycosylation in IgAN may be restricted to a small fraction of B-cell populations rather than being a uniform property of all IgA1-producing cells [52, 53]. Similarly, Gd-IgA1-secreting cells in the peripheral blood of patients with IgAN are enriched for the plasmablast-like cells with mucosal homing receptors [54].

These findings must be interpreted within the context of the model system. The facts that primary PBMCs typically do not survive longer than 7-12 days in culture and contain only a small amount of IgA1^+^ cells limit our investigations to mechanistic studies in immortalized cells. In EBV-immortalized B cells, the signaling apparatus depends on constitutive activation of the p65 NF-κB pathway, as seen in control cells showing phosphorylation of S536 (**Fig. 1F)**. Despite the constitutive activation, we observed significant distribution of production of Gd-IgA1 and IgA1 between cells from different donors, suggesting that differences in critical signaling mechanisms controlling glycosyltransferase activity are not solely dependent on p65 NF-κB canonical activation (**Figs. 1C, D**). Additionally, EBV immortalization has been shown to not alter the glycosylation of IgA1, as the phenotypes of Gd-IgA1 produced by immortalized cells are comparable to that of IgA1 in donor serum [23]. Constitutive p65 NF-κB activation may explain why inflammatory induction via cytokine stimulation using IL-6-mediated activation of STAT3 increases production of Gd-IgA1, whereas the same affect is mediated by inhibition of p65 NF-κB [31]. TPCA-1 is known to modestly inhibit activation of STAT3. We confirmed this effect (**Supp Fig. 3D**) but, in theory, inhibition of STAT3 would be expected to decrease production of Gd-IgA1. In a prior publication, we found no effect of STAT3 inhibition on production of Gd-IgA1 without stimulation by IL-6 [31]. When we exposed immortalized B cells to TPCA-1, we elicited a short-lived decrease in p65 S536 phosphorylation at 1 hr, that reversed over 24 hrs. This transient nature of p65 NF-κB inhibition highlights the sustained changes in phenotype we observed after 48 hr, including the SSC population shift and decrease in SP1^+^ population frequency, that could represent downstream consequences persisting beyond acute disruption.

EBV-immortalized B-cell populations are not homogenous but instead show substantial subpopulation diversity. TPCA-1 induced changes in cell morphology manifested by a shift from SSC-low to SSC-high populations, with corresponding greater cell-surface HPA binding and CD138 expression (**Fig. 3, Suppl. Fig. 2**). These findings are consistent with a plasma-cell-like phenotype and agree with other studies showing cell-surface glycophenotyping can be used to assess stages of B-cell maturation [55]. Notably, Gd-IgA1-producing cells are enriched for CD138 in the circulation of patients with IgAN [54]. Immunophenotyping flow analysis at baseline showed a variety of subpopulations across 15 different markers but, as these are EBV-immortalized cells, they do not fit perfectly into traditional B-cell subpopulation “buckets”. This feature inherently limits use of EBV-immortalized cells, as viral transformation alters normal cell signaling. Regardless, the SSC-based groupings revealed subpopulations with distinct intracellular signaling profiles. Intracellular analysis showed substantially lower SP1 MFI, SP1 nuclear translocation, and IgA MFI in the SSC-high subpopulations with a trend for lower p65 NF-κB MFI, indicating these cells may be critical Gd-IgA1 producers (**Figs. 4, 5**). The lower SP1 MFI and SP1 nuclear translocation in SSC-high cells **(Figs. 4C, 5B**, respectively) are consistent with reduced *C1GALT1* translational support, which helps to explain the increased Gd-IgA1 production. Furthermore, unsupervised clustering analysis found that the most significant correlation between the flow parameters and Gd-IgA1 production was the SP1 subpopulations (**Fig. 6**). Combining the unsupervised analysis with hypothesis-driven SP1 analyses strengthens the concept that SP1 plays a mechanistically relevant role in Gd-IgA1 production. Taken together, these data suggest a model in which acute IKKβ/p65 NF-κB inhibition leads to increased frequency of SSC-high/SP1^-^ subpopulations that are likely responsible for Gd-IgA1 production.

The ability of TPCA to arrest cellular mitosis did not reach significance (**Fig. 1A**; p = 0.35), but if we dropped the outlier, it did reach significance (p = 0.02). This raises the question whether the primary role of NF-κB in Gd-IgA1 secreting B cells is inflammatory signaling or whether it is related to its control of cell cycle. Multiple studies have shown that in PBMCs from IgAN patients, IgA^+^ B cells and plasma cells constitute a higher percent of total cells than in healthy controls and this higher frequency correlates with elevated proteinuria [56]. TPCA-1 inhibited p65 phosphorylation at S536 at 1 hr (**Fig. 1F-I)**, and yet total p65 NF-κB protein levels, population frequency, and nuclear translocation were not significantly altered at 48 hr across multiple platforms. These findings indicate that long-term phenotypic effects of IKKβ inhibition do not depend on continuous suppression of p65 protein, but perhaps initiate downstream events that then persist independently. Our data suggest a complex regulatory mechanism for controlling glycosyltransferases as an interplay of inflammatory and growth-related transcription factors that remove the normal homeostatic control of these enzymes. Future studies examining progression of the cell cycle, maturation state, and glycosylation may be critical for understanding how these processes influence autoantigen production.

Our study opens several avenues for further investigation. ChiP Seq and Co-IP assays would determine whether SP1 and p65 directly co-occupy the *C1GALT1* promoter in IgA1-producing B cells. SP1 knockdown and/or overexpression studies would assess whether SP1 is sufficient to regulate *C1GALT1* expression and Gd-IgA1 production independently of NF-κB. Intermediate timepoints between 1 and 48 hr would clarify the kinetics of the proposed NF-κB/SP1/C1GalT1 cascade. Validation of our findings in primary B cells from mucosal tissue would strengthen the translational relevance of these effects. Incorporating more specific markers of plasmablast lineage with the addition of glycophenotyping could more precisely characterize Gd-IgA1-producing cells.

Herein we provide the first experimental evidence that canonical p65 NF-κB signaling may play a role in the regulation of IgA1 *O*-glycosylation in human B cells. Our findings link the GWAS-identified susceptibility loci related to NF-κB pathways to the production of the Gd-IgA1, the autoantigen in the pathogenesis of IgAN. This signaling process likely involves SP1-dependent transcriptional control of *C1GALT1*, where loss of SP1 enhances Gd-IgA1 production. We anticipate that the involvement of growth factors needs to be considered, as studies in cancer have shown their role in controlling *C1GALT1* expression [57, 58]. Our findings also confirm that galactose-deficient glycophenotypes are not ubiquitous among IgA1-producing cells. However, these forms may be enriched within a plasmablast-like subpopulation characterized by reduced SP1 nuclear activity, such that control of B-cell differentiation and glycosylation are important for production of Gd-IgA1. Understanding how specific signaling axes govern IgA1 glycosylation will be essential for understanding the autoimmune molecular origins of IgAN, leading to more pathogenesis-targeted therapies for patients.

## Credit authorship contribution statement

**Taylor Person:** Writing – review & editing, Writing – original draft, Methodology, Investigation, Formal analysis, Data curation, Visualization, Validation, Conceptualization. **Maggie Phillips:** Investigation, Formal analysis. **Terri Rice:** Investigation, Methodology. **Stacy Hall:** Resources, Methodology. **Bruce A. Julian:** Resources, Conceptualization. **Dana V. Rizk:** Project Administration, Resources, Conceptualization. **Jan Novak:** Writing – review & editing, Methodology, Data curation, Visualization, Conceptualization, Funding acquisition, Resources, Supervision, Project Administration. **Colin Reily:** Writing – review & editing, Methodology, Data curation, Visualization, Conceptualization, Formal analysis, Data curation, Validation, Funding acquisition, Resources, Supervision, Project Administration.

## Acknowledgements

We thank Sagar Hanumanthu at the UAB CFCC Flow Cytometry Core for all his assistance and guidance with instrumentation and compensation. We thank Jamie Clemons for structuring the profiles and demographics of the donors of the peripheral blood mononuclear cells that were used to develop the immortalized B-cell lines.

