## Supplementary figures and tables for "Inhibition of p65 NF-κB enhances production of galactose-deficient IgA1 through suppression of *C1GALT1* and SP1 in plasmablast-like cell subpopulations"

### Supplementary figures and table

**Supplementary Data**

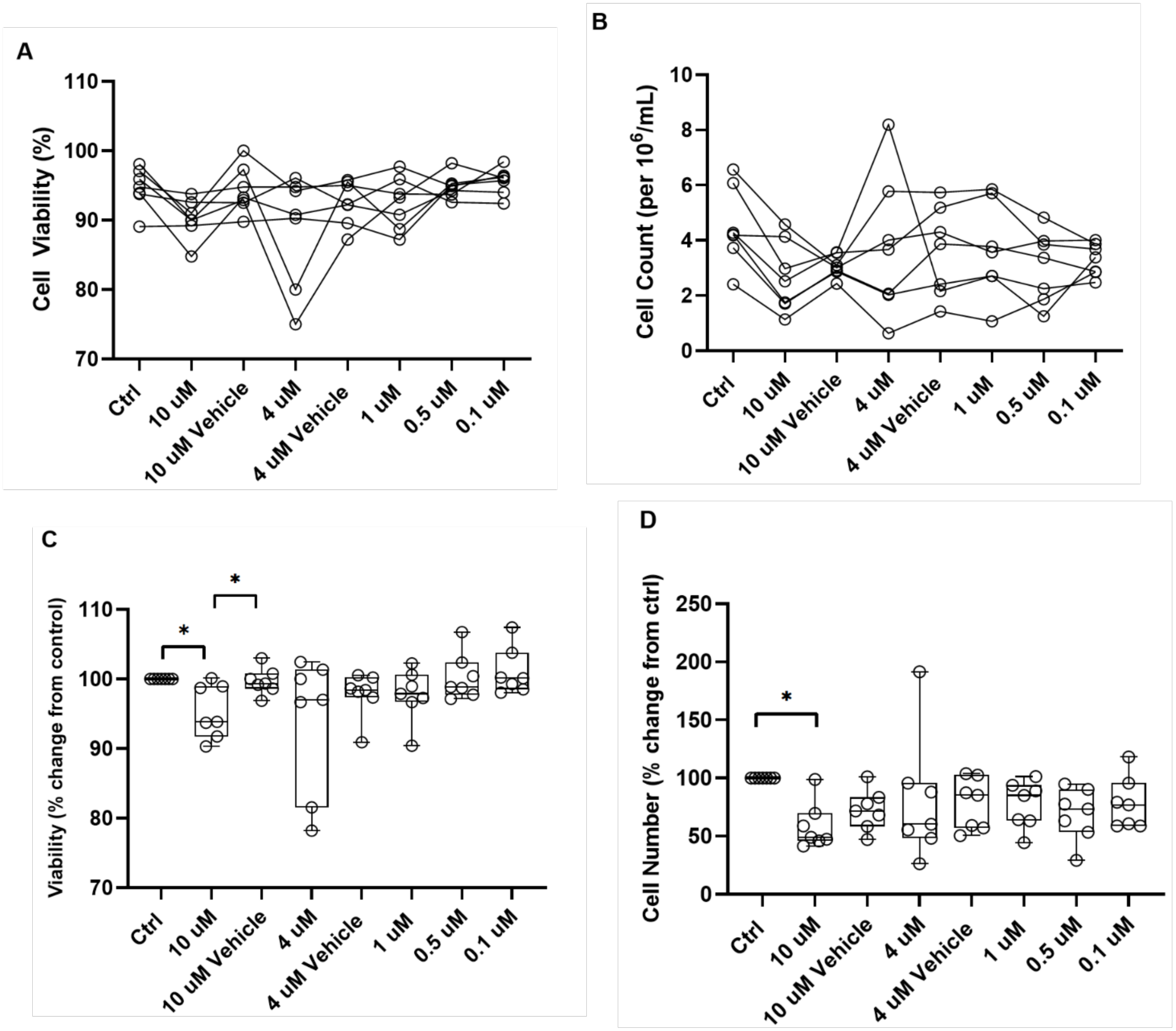

**Supplementary Data Figure 1: Dose response for TPCA-1 on EBV-immortalized B cells.** Cells were plated at $1x10^6$/mL. **A)** Cell viability expressed as percent of sample after 48 hr. **B)** Cell count ($x10^6$) after 48 hr. **C)** Cell viability, percent change from control after 48 hr. **D)** Cell number, percent change from control after 48 hrs. Vehicle is DMSO at the same concentration as the respective TPCA-1 treatment. Ctrl = control. ***p = 0.01.**

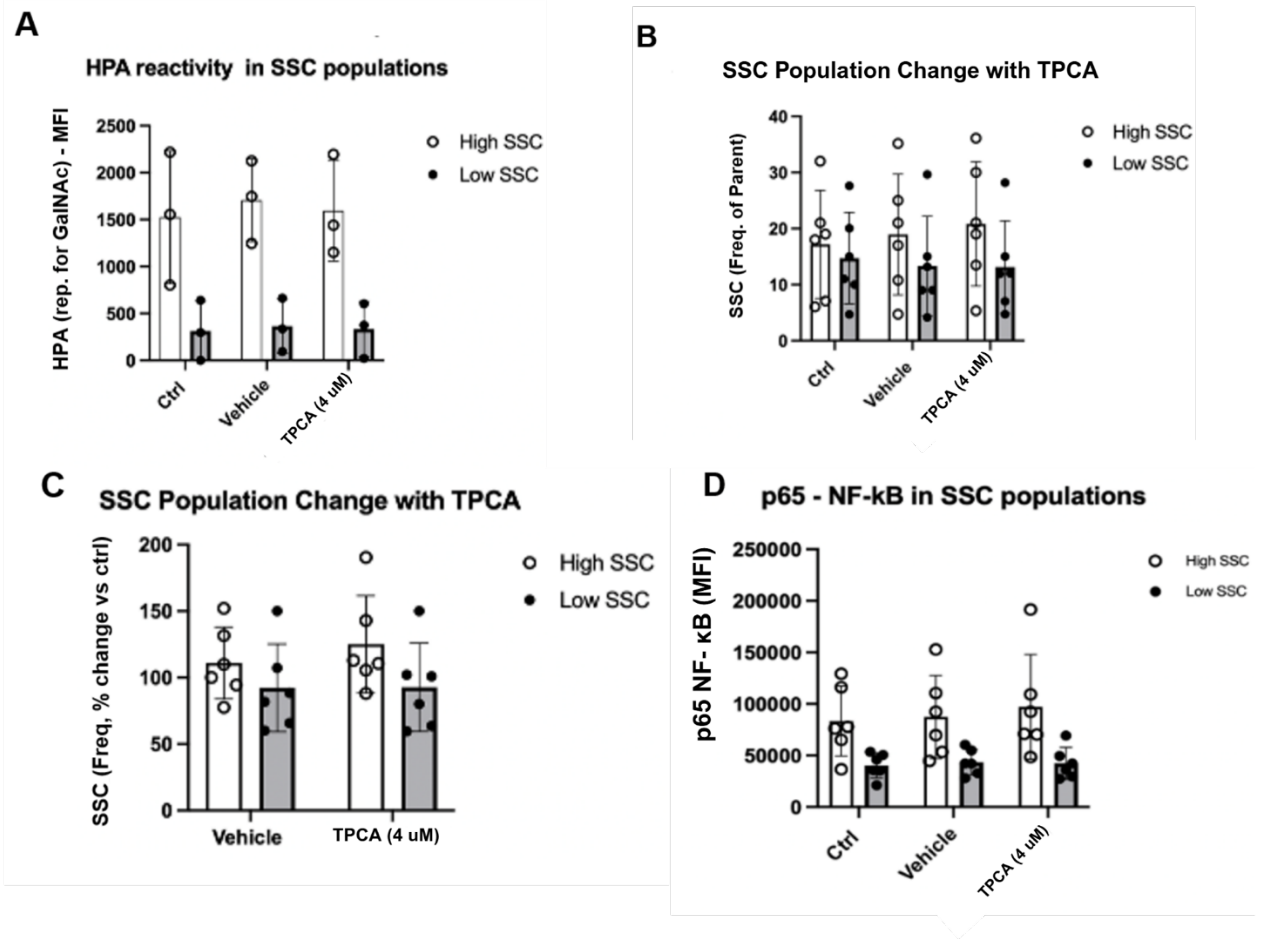


**Supplementary Data Figure 2: Characterization of SSC subpopulations at 1 hr. A)** HPA reactivity (lectin specific for terminal GalNAc) in high and low SSC-A subpopulations across control, vehicle, and TPCA-1 conditions. **B)** Frequency of high and low SSC-A subpopulations as percent gated from parent across treatment conditions. **C)** Percent change from control in high and low SSC-A subpopulations after vehicle or TPCA-1 treatment. **D)** p65 NF-κB (S536) mean fluorescence intensity in high and low SSC-A subpopulations across treatment conditions. Data points represent individual samples. Bars represent mean +/- SD.

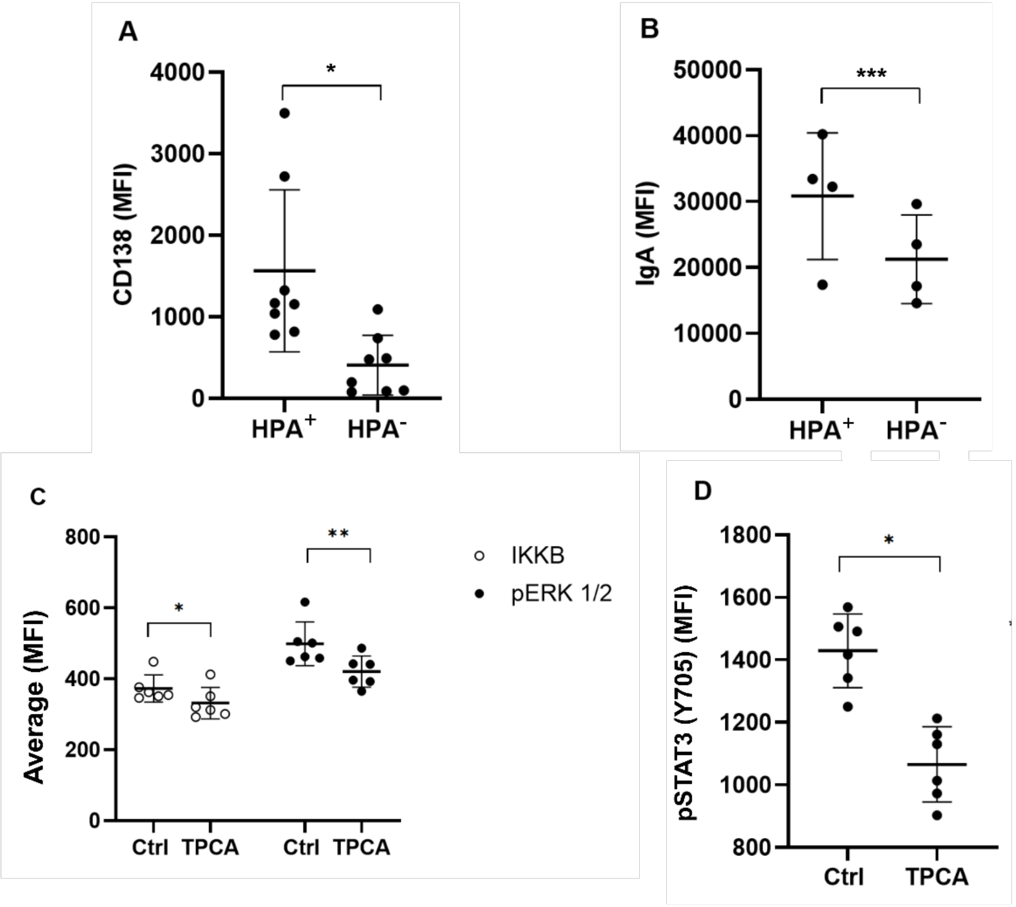


**Supplementary Data Figure 3: Baseline association of HPA$^+$ subpopulations and effects of TPCA on IKKβ, pERK, and pSTAT3.** **A)** CD138 median fluorescence intensity in HPA$^+$ and HPA$^-$ cells ($p<0.01$). **B)** IgA median fluorescence intensity in HPA$^+$ and HPA$^-$ cells ($p=0.05$). **C)** IKKβ (left) and pERK 1/2 (right) average median fluorescence intensity (MFI) in control and TPCA-1-treated cells at 1 hr. TPCA-1 decreased IKKβ ($p<0.01$) and increased pERK 1/2 ($p=0.02$). **D)** pSTAT3 (Y705) average MFI in control and TPCA treated cells at 1 hr ($p<0.01$). Bars represent mean +/- SD. *= $p < 0.01$. **= $p < 0.02$. ***= $p < 0.05$.

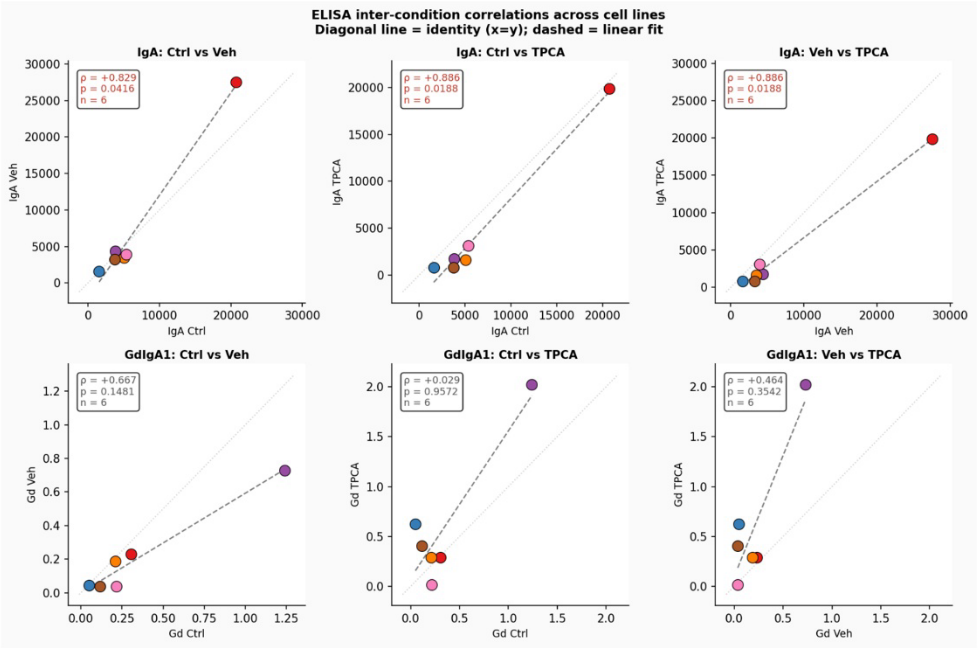


**Supplementary Data Figure 4: IgA1 and Gd-IgA1 inter-condition correlations across cell lines.** IgA1 production (top row) and Gd-IgA1 production (bottom row) were compared across control, vehicle, and TPCA-1 conditions. Each data point represents an individual donor. Dashed line represents linear fit. Spearman correlation coefficients (ρ) and p-values are shown for each comparison. IgA1 production was significantly correlated across all condition pairs (top row), while Gd-IgA1 production showed no significant correlation between any condition (bottom row). IgA units (ng/mL), Gd-IgA1 units expressed as rGd-IgA1standard (U).

**Supplementary Table 1.** Demographic and baseline clinical characteristics of IgA nephropathy patients and healthy control donors.

| Donor ID | Sex | Age (yr) | Ethnicity | SCr (mg/dL) | eGFR (mL/min/1.73 m²) | UPCR (g/g) |
|---|---|---|---|---|---|---|
| IgAN1 | Male | 34 | Non-Hispanic | 1.9 | 47 | 0.337 |
| IgAN2 | Male | 23 | Non-Hispanic | 0.8 | 128 | 0.492 |
| IgAN3 | Male | 33 | Non-Hispanic | 3.5 | 23 | 3.596 |
| IgAN4 | Male | 35 | Non-Hispanic | 3.7 | 21 | 1.492 |
| IgAN5 | Male | 63 | Non-Hispanic | 4.1 | 16 | 2.888 |
| IgAN6 | Male | 43 | Non-Hispanic | 4.9 | 14 | 1.451 |
| IgAN7 | Male | 46 | Non-Hispanic | 1.5 | 58 | NA (UP<6) |
| IgAN8 | Male | 33 | Non-Hispanic | 1 | 102 | 0.048 |
| IgAN9 | Male | 43 | Non-Hispanic | 1.1 | 85 | 0.044 |
| IgAN10 | Female | 47 | Non-Hispanic | 0.9 | 79 | 0.606 |
| IgAN11 | Female | 35 | Non-Hispanic | 14.3 | 3 | 0.805 |
| IgAN12 | Male | 73 | Non-Hispanic | 1.3 | 58 | 0.309 |
| IgAN13 | Female | 56 | Non-Hispanic | 8.3 | 5 | 1.924 |

| Donor ID | Sex | Age (yr) | Ethnicity | SCr (mg/dL) | eGFR (mL/min/1.73 m²) | UPCR (g/g) |
|---|---|---|---|---|---|---|
| IgAN14 | Male | 54 | Non-Hispanic | 1.2 | 72 | 0.641 |
| IgAN15 | Female | 41 | Non-Hispanic | 0.6 | 116 | 1.043 |
| IgAN16 | Male | 44 | Non-Hispanic | 1.4 | 64 | 0.051 |
| IgAN17 | Male | 37 | Non-Hispanic | 1.1 | 89 | 1.76 |
| IgAN18 | Male | 38 | Non-Hispanic | 1.5 | 61 | 2.5 |
| IgAN19 | Male | 41 | Non-Hispanic | 1.3 | 71 | 0.108 |
| HC1 | Male | 72 | Non-Hispanic | 0.5 | 108 | 0.388 |
| HC2 | Male | 46 | Non-Hispanic | 1 | 94 | 0.593 |
| HC3 | Female | 63 | Non-Hispanic | 1.3 | 46 | ND |
| HC4 | Female | 42 | Non-Hispanic | 1 | 72 | 0.038 |
| HC5 | Male | 61 | Non-Hispanic | 1 | 86 | NA (UP<6) |
| HC6 | Male | 36 | Non-Hispanic | ND | ND | 0.49 |
| HC7 | Female | 37 | Non-Hispanic | 1.1 | 66 | NA (UP<6) |
| HC8 | Male | 17 | Non-Hispanic | 0.9 | 128 | 0.012 |

| Donor ID | Sex | Age (yr) | Ethnicity | SCr (mg/dL) | eGFR (mL/min/1.73 $m^2$) | UPCR (g/g) |
|---|---|---|---|---|---|---|
| HC9 | Male | 15 | Non-Hispanic | ND | ND | 0.014 |
| HC10 | Male | 41 | Non-Hispanic | 1.1 | 86 | 0.376 |
| HC11 | Male | 47 | Non-Hispanic | 0.9 | 106 | 0.666 |
| HC12 | Male | 46 | Non-Hispanic | 1.2 | 76 | NA (UP<6) |

***Abbreviations:*** IgAN, IgA nephropathy; HC, healthy control; SCr, serum creatinine; eGFR, estimated glomerular filtration rate (2021 CKD-EPI equation, without race coefficient); UPCR, urine protein-to-creatinine ratio.

**Supplementary Table 2:** Flow cytometry reagents used in the methods.

| Reagent/Antibody | Target | Conjugate | Catalog # | Manufacturer | Dilution |
|---|---|---|---|---|---|
| HPA lectin | Terminal GalNAc (cell surface) | Alexa Fluor 647 | L32454 | Invitrogen | 1:100 |
| Phospho-NF-κB p65 (S536) monoclonal antibody | p65 NF-κB (phospho-S536) | Unconj. | MA5-15160 (T.849.2) | Invitrogen | 1:1000 |
| NF-κB p65 whole protein recombinant rabbit monoclonal antibody | p65 NF-κB (total protein) | Unconj. | 701079 (4-2H22L23) | Thermo Fisher | 1:1000 |
| Goat anti-rabbit IgG (H+L) | Secondary (rabbit primary) | AF488 | 4050-30-8 | Southern Biotech | 1:1000 |
| SP1 rabbit monoclonal antibody | SP1 | Unconj. | 9389S | Cell Signaling | 1:1000 |
| Donkey anti-rabbit IgG | Secondary (for SP1) | AlexaFluor-647 | 406414 | BioLegend | 1:1000 |
| DAPI (4',6-Diamidino-2-Phenylindole, Dilactate) | Nuclear stain | n/a | 50-403-592 | ThermoFisher | 1:1000 |
| Mouse anti-human IgA PE | Cell surface and total IgA1 | PE | OB9130-09 | Southern Biotech | 1:500 |
| CD138 | CD138 (Syndecan-1) | PE-Cy7 | 25-1389-42 | Invitrogen | 1:100 |
